# Beyond point estimates: quantifying predictive uncertainty reveals hidden dimensions of biological age acceleration and improves risk interpretation

**DOI:** 10.64898/2026.08.05.742349

**Authors:** Chen Wang, Hanqing Wu, Shinichi Namba, Jun Young Park, the BioBank Japan Project, Koichi Matsuda, Yukinori Okada, Zihuai He, Iuliana Ionita-Laza

## Abstract

Biological age estimates are increasingly used to study aging, disease risk, and mortality, yet their predictive uncertainty is rarely quantified. Consequently, conventional age-gap measures can treat deviations as equally informative even when the underlying biological age predictions differ substantially in reliability. We developed a framework for uncertainty-aware biological aging that generates calibrated prediction intervals and individualized probabilities of accelerated or decelerated aging alongside point estimates. We applied this framework to the UK Biobank Pharma Proteomics Project, evaluating three composite and eleven organ-specific biological age clocks. Predictive uncertainty varied substantially both within and across clocks, revealing that apparently extreme age gaps can differ markedly in the strength of evidence supporting accelerated or decelerated aging. In particular, low-accuracy clocks, including many organ-specific clocks, provided little evidence for confidently accelerated or decelerated aging. Beyond biological age gaps, prediction-interval width was independently associated with disease risk and mortality, particularly for composite, brain, and immune clocks, suggesting that predictive uncertainty captures an additional dimension of biological aging that may reflect increased molecular heterogeneity and dysregulation associated with aging and disease. We replicated these findings in Biobank Japan and an independent clinical cohort from Stanford. By incorporating individual-specific predictive uncertainty, our framework provides a more informative characterization of biological aging and enables improved individual-level risk stratification for disease prevention and longitudinal monitoring.

## Introduction

Biological aging is a multifaceted process that varies considerably among individuals of the same chronological age, leading to differences in functional capacity, disease risk, and mortality. This heterogeneity has motivated the development of biological clocks, i.e., computational models that estimate biological age from molecular and physiological biomarkers. Advances in high-throughput omics technologies have enabled aging clocks based on DNA methylation, transcriptomic, proteomic, metabolomic, and imaging-derived data (Horvath 2013; Horvath and Raj 2018; Tanaka et al. 2018; Lehallier et al. 2019; Rutledge et al. 2022). These clocks have demonstrated that machine learning models trained on various molecular profiles can accurately predict chronological age while capturing variation associated with health outcomes.

A central concept in biological clock research is the distinction between predicted biological age and chronological age. The residual difference between these quantities, often termed “age gap”, is interpreted as a measure of accelerated or decelerated aging. Positive age gaps have been associated with increased risk of chronic disease, frailty, cognitive decline, and mortality, whereas negative age gaps may reflect resilience or healthy aging (Rutledge et al. 2022; Argentieri et al. 2024; Y. Wang et al. 2026). However, aging does not occur uniformly across the body. Different organs and physiological systems age at different rates depending on genetic predisposition, environmental exposure, disease burden, and lifestyle factors. Consequently, organismal aging is increasingly understood as a heterogeneous and organ-specific phenomenon rather than a single global process (Tian et al. 2023).

This realization has led to the development of organ-specific biological clocks (D. H. Oh et al. 2023; H. S.-H. Oh et al. 2025; Y. Wang et al. 2026). Instead of predicting a single whole-body biological age, these models estimate the biological age of individual organs or systems such as the brain, heart, liver, kidney, immune system, or musculoskeletal system. Organ-specific clocks are typically trained using biomarkers most relevant to the target tissue, including imaging features, circulating proteins, transcriptomic signatures, or clinical laboratory measures. For example, brain age models derived from magnetic resonance imaging have shown associations with neurodegenerative disease and cognitive impairment (Cumplido-Mayoral et al. 2025).

While biological age prediction models are often summarized by point estimates, uncertainty quantification can provide critical additional information for interpreting individual age gaps. This is particularly relevant for organ-specific clocks, where differences in the amount of biomarker information available across organs may lead to varying levels of prediction precision. When molecular profiles contain limited age-related information, age predictions regress toward the population mean, resulting in age gaps that carry little individual-level information. As predictive accuracy improves, deviations between predicted and chronological age increasingly reflect an individual’s departure from age-specific molecular expectations, rather than inaccuracies in the predictor. We therefore use quantile regression (Koenker et al. 1978; C. Wang, T. Wang, et al. 2024; F. Wang et al. 2025; C. Wang, F. Wang, et al. 2025; Wu et al. 2026) to estimate the conditional distribution of age given an individual’s molecular profile, enabling probabilistic assessment of whether an individual’s predicted age deviates from age-specific expectations rather than relying solely on point estimates. The proposed framework produces calibrated prediction intervals for biological age and individualized probabilities of accelerated or decelerated aging, providing a probabilistic characterization of biological aging with distinct advantages over the traditional point estimate-based approaches commonly used in the biological aging literature.

Building on this probabilistic formulation, we contrast our approach with the usual practice in biological aging studies of identifying accelerated or decelerated agers using global thresholds on age gap distributions (e.g., deviations exceeding a fixed number of standard deviations from the population mean). By modeling the conditional distribution of age given an individual’s molecular profile, quantile regression induces a personalized reference distribution, enabling outlier status to be assessed in terms of individualized tail probabilities rather than population-level cutoffs. As a result, individuals with identical age gaps from the same biological age clock need not provide the same evidence for accelerated or decelerated aging, as their classification depends on their position within the covariate-dependent uncertainty landscape.

To illustrate this distinction between traditional age gap analyses and a conditional quantile regression framework, consider two individuals with the same estimated biological age gap of +10 years. Under the standard approach, both individuals would be classified as equally accelerated agers. However, the precision with which age can be inferred from their molecular profiles may differ substantially. For one individual, the conditional age distribution implied by their molecular profile may be narrow, making a 10-year discrepancy highly unusual. For the other, the corresponding distribution may be much broader, such that a 10-year discrepancy falls well within the range expected among individuals with similar molecular characteristics. Quantile regression distinguishes between these scenarios by evaluating age gaps *relative* to the distribution of age gap conditional on an individual’s molecular profile rather than a single population-wide age-gap distribution. Consequently, individuals with identical age gaps may differ substantially in the strength of evidence for age-molecular profile discordance.

An important yet largely unexplored dimension of biological aging is the uncertainty with which biological age can be inferred from molecular profiles. We propose that this uncertainty is not merely a statistical limitation of biological age estimation, but may itself represent a complementary biomarker of aging. Variation in predictive uncertainty may capture differences in the heterogeneity, coherence, or dysregulation of molecular processes underlying aging, providing information that is not reflected in the magnitude of the biological age gap alone. By quantifying individual-specific uncertainty alongside biological age, our framework captures two complementary dimensions of biological aging: the extent to which an individual’s molecular profile indicates an older or younger biological state, and the degree to which that state can be inferred with confidence.

We first present an overview of the uncertainty quantification framework, followed by primary results from the UK Biobank Pharma Proteomics Project (UKB-PPP) and replication analyses in Biobank Japan and a pooled Stanford cohort comprising samples from the Stanford Alzheimer’s Disease Research Center (ADRC) and the Stanford Aging and Memory Study (SAMS).

## Results

### Overview of the study

To predict organ-specific biological age we first used plasma proteomics data from the UK Biobank Pharma Proteomics Project, part of the UK Biobank data (Sun et al. 2023; *n* = 44, 498 individuals, *p* = 2, 916 proteins), as employed in previous studies (H. S.-H. Oh et al. 2025). We fitted penalized quantile regression models using FS-QRPPA (Methods) (Wu et al. 2026), a scalable optimization algorithm, to predict biological age at several quantile levels, including *τ* = 0.025, 0.1, 0.2, …, 0.8, 0.9, 0.975. Specifically, we modeled the conditional distribution of chronological age *y*_*i*_ given proteomic features ***x***_*i*_ using quantile regression, yielding estimates 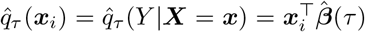, where *τ* is a quantile level and 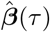 was fitted on the training set using FS-QRPPA. For 0 < *τ* < 0.5, the prediction interval PI_1−2*τ*_ with nominal coverage level 1 − 2*τ* for chronological age given the proteomic profile ***x***_*i*_ is then given by

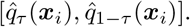

Unlike a confidence interval, which reflects uncertainty in estimating a model parameter, a prediction interval captures the conditional variability of biological age given an individual’s molecular profile, thereby reflecting both the expected range of outcomes and the heterogeneity in their predictability.

Following the common convention in the aging-clock literature that the predicted chronological age can serve as a measure of biological age (Rutledge et al. 2022), we define the biological age score as the median prediction 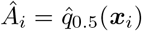. Let *G*_*i*_ denote the raw age gap for individual *i*. A point estimate of the raw age gap is 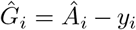, where *y*_*i*_ is the observed chronological age. However, the raw age gap is typically negatively correlated with chronological age due to regression-to-the-mean effects, whereby younger individuals tend to have overestimated ages and older individuals tend to have underestimated ages. To address this, we instead focused on the adjusted age gap 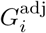, which removes the chronological-age-dependent trend by subtracting the conditional mean prediction at chronological age *y*_*i*_ (Methods). In the same spirit, we constructed prediction intervals PI_1−2*τ*_ for 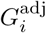.

Furthermore, we reported the estimated upper-tail probability

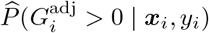

as a measure of accelerated aging relative to individuals of the same chronological age.

We divided the dataset into three separate sets, including training (50%), validation (10%) and test (40%). For each individual in the test data we predicted its biological age, adjusted age gap along with prediction intervals, and tail probabilities as described above. We trained several prediction models, including:

1. Composite models:
  - Conventional: based on all proteins.
  - Multiorgan: based on all organ-enriched proteins across all organs.
  - Organismal: based on all remaining proteins after excluding organ-enriched proteins.
2. Organ-specific models: based on organ-enriched proteins from a single organ (artery, brain, etc.), as defined by previous studies (D. H. Oh et al. 2023; H. S.-H. Oh et al. 2025; Table S1).

We compared our measure with the standardized age gap commonly used in the biological aging literature. Specifically, we first estimated the adjusted age gap 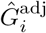, and then standardized it by subtracting its sample mean and dividing by its sample standard deviation. Following usual practice, we refer to these standardized residuals as standardized age gaps. We perform replication analyses in Biobank Japan and the Stanford ADRC and SAMS combined cohort (Methods). A graphical overview of our study is provided in Figure 1.

**Figure 1:**
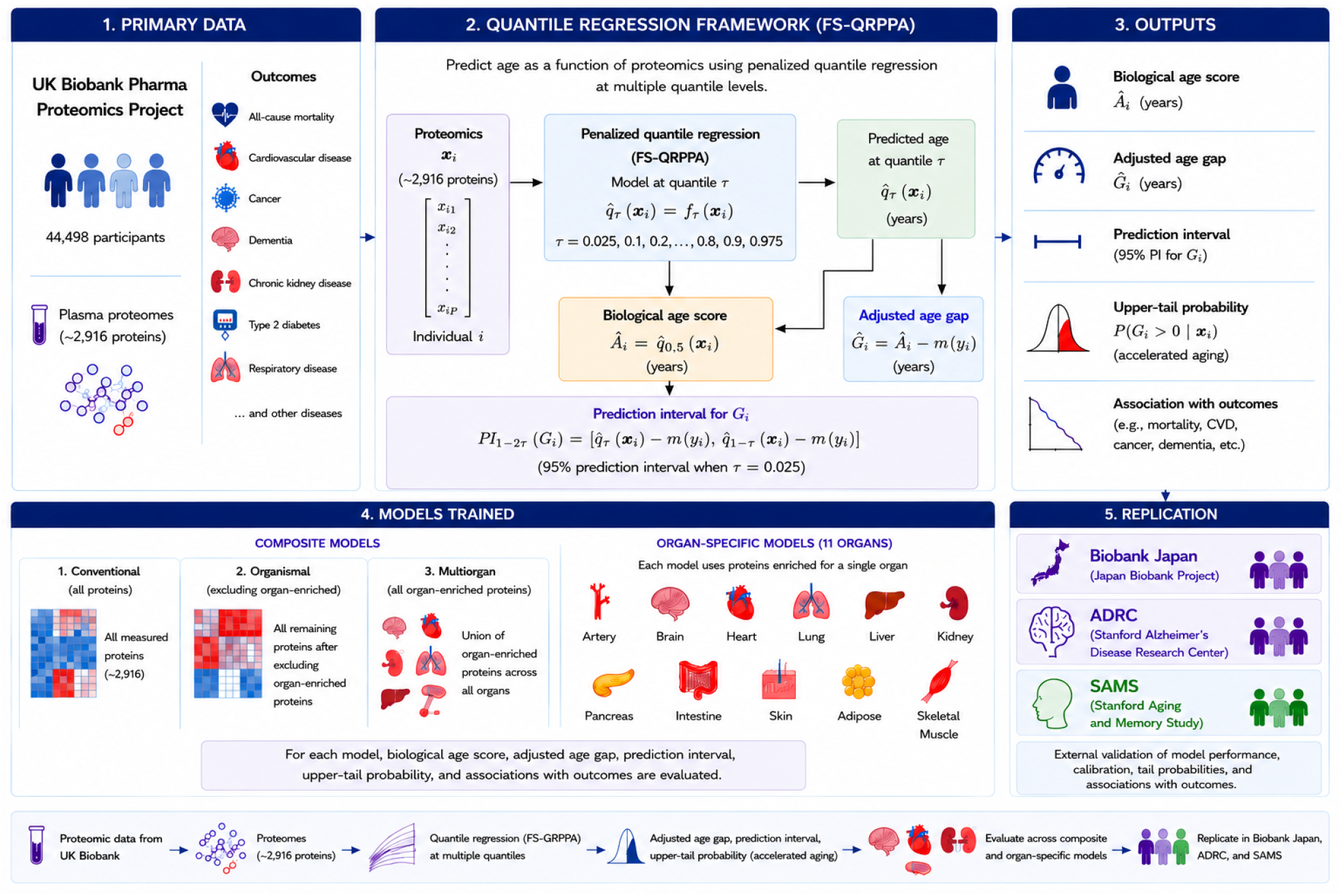
Workflow of FS-QRPPA framework for proteomics-based biological age prediction.

### Organ age prediction accuracy scales with protein availability

We first evaluate age prediction accuracy across models (Figure 2(a)). Composite models, such as the conventional, organismal and multiorgan models, yield biological age predictions that are highly correlated with chronological age (Pearson *r* ≈ 0.9) and exhibit low predictive error. In contrast, organ-specific models based on organ-enriched proteins show weaker age correlations, which track closely with the number of organ-specific proteins used in prediction. This pattern is particularly pronounced for kidney, muscle, pancreas, lung, adipose tissue, and heart, which exhibit markedly weaker correlations and higher prediction errors, potentially reflecting limited protein coverage in model training. Consistent with these findings, correlations across organs are also lowest for these tissues for both biological age predictions and adjusted age gaps (Figure S1).

**Figure 2:**
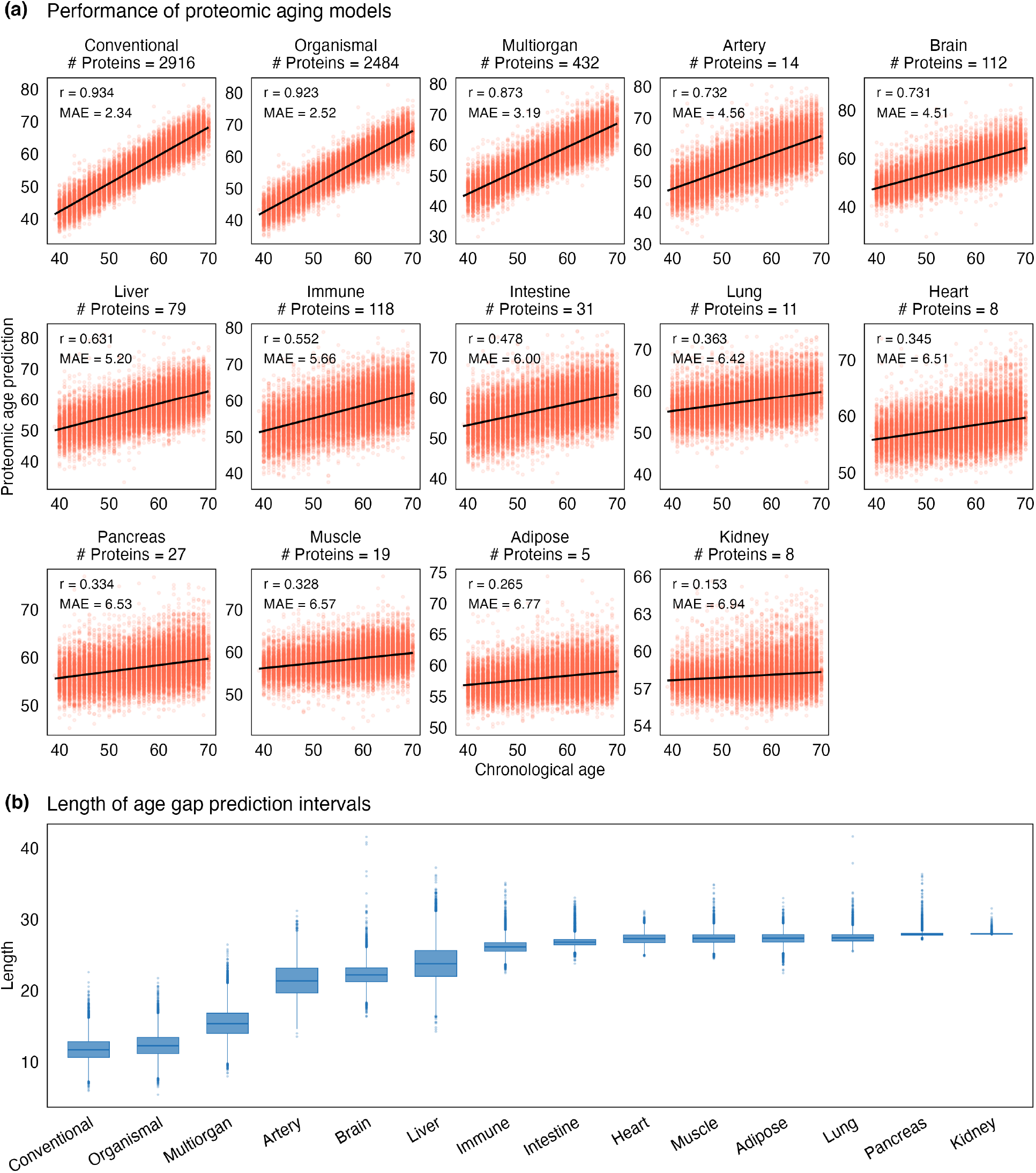
Clock accuracy and prediction interval length in UK biobank. (a) Performance of proteomic aging models across different clocks. Each panel includes the number of protein features used to train the model, the Pearson correlation coefficient (*r*) between the median prediction and chronological age, and the Mean Absolute Error (MAE) measured in years. (b) Distribution of the prediction interval lengths for different organ-specific and composite clocks in UK Biobank.

### Uncertainty quantification via prediction intervals and tail probabilities

As illustrated above, there is substantial variation in the accuracy and uncertainty of biological age predictions across different organs. Even for the same organ model, uncertainty can vary a lot from person-to-person. To formally quantify this uncertainty, we construct prediction intervals that characterize the spread of the conditional predictive distribution by providing the lower and upper bounds within which the true chronological age is expected to lie with a specified probability. We take *τ* = 0.025 and 0.975 unless specified otherwise, yielding 95% prediction intervals. We evaluate empirical coverage as the proportion of individuals for whom the true chronological age falls within the interval. Overall, both composite and organ-specific models achieve coverage close to the nominal 95% level (Figure S2). Since the adjusted age gap is obtained from the raw age gap via a location-shift transformation, the resulting prediction intervals are translation-invariant, and their coverage is therefore identical to that of the underlying conditional quantile model above.

The resulting intervals are adaptive to the data. In particular, they are asymmetric around the age gap estimate and can vary in length from person to person (Figure 3). Furthermore, prediction interval lengths vary substantially across organ-specific biological age models, indicating marked differences in prediction precision and reliability between organs (Figures 2(b) and 3). Composite clocks, including organismal, conventional and multiorgan clocks, have the narrowest intervals consistent with their high prediction accuracy, followed by brain, arterial and immune clocks. In contrast, kidney, muscle, adipose, lung, pancreas and heart clocks showed substantially greater uncertainty. This greater uncertainty observed for some organs may reflect reduced predictive information, including smaller protein feature sets and weaker model performance, which is not captured by point estimates alone. We also find that prediction uncertainty varies substantially across individuals for the same biological age model (Figure 2(b)). Less accurate clocks such as heart, lung, pancreas, kidney tend to produce uniformly broad prediction intervals, whereas more accurate clocks such as composite, immune, artery and brain often exhibit pronounced heterogeneity in uncertainty across individual molecular profiles.

**Figure 3:**
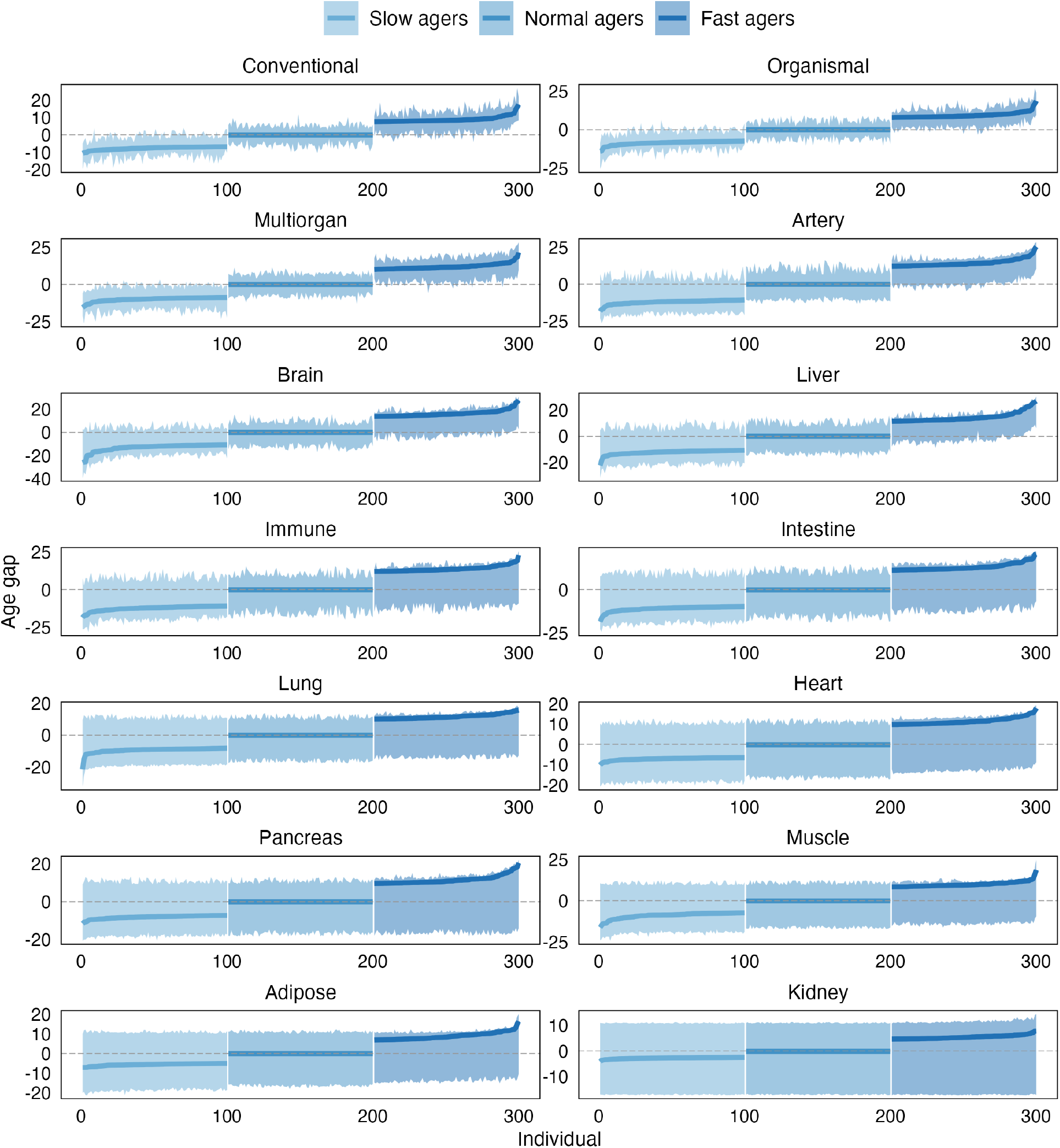
Adaptive prediction intervals across different aging clocks and organs in UK Biobank. For each individual on the x-axis, the adjusted age gaps (solid blue points) along with their corresponding prediction intervals are shown. Three groups of 100 individuals each are shown: slow agers (left), normal agers (middle), and fast agers (right). The prediction intervals are adaptive: they vary in length from person to person, and can be highly asymmetric around the central prediction line.

Prediction intervals quantify uncertainty in biological age estimates but may provide limited granularity for assessing individual-level deviations from chronological age when uncertainty is substantial. Tail probabilities therefore offer a more sensitive measure of evidence for accelerated or decelerated aging, enabling individuals to be ranked along a continuous spectrum of age deviation even when their prediction intervals include chronological age. Our framework integrates these perspectives by reporting, for each individual, point predictions of adjusted age gap, calibrated prediction intervals, and tail probabilities. Although tail probabilities are highly correlated with standardized age gaps across models (Figure S3(a)), they provide an important complementary dimension by explicitly incorporating predictive uncertainty. This probabilistic framework distinguishes individuals whose apparent accelerated or decelerated aging is strongly supported by their molecular profiles from those whose extreme age gaps may instead reflect substantial prediction uncertainty, enabling more reliable identification of truly accelerated or decelerated aging profiles, as illustrated below. Notably, for the most accurate aging models, both standardized age gaps and tail probabilities exhibit only weak correlations with prediction interval length (Figure S3(b–c)), indicating that the magnitude of biological age deviation and the certainty with which it can be inferred represent largely distinct aspects of biological aging.

For individuals that may be classified as outliers based on threshold-based criteria, such as those with a standardized age gap greater than 1.5 or less than − 1.5 in a given aging model, the tail probabilities behave as expected (Figure 4). Specifically, for accurate clocks such as conventional, organismal, multiorgan, artery, immune and brain models, tail probabilities are concentrated near 1.0 or 0.0, indicating that indeed these individuals lie in the extreme tails of the estimated conditional age distributions. In contrast, for lower-accuracy clocks such as kidney, adipose, and heart, tail probabilities are substantially less decisive, with many individuals exhibiting moderate values despite similarly large standardized age gaps (Figure 4). This pattern reflects the greater uncertainty and broader conditional age distributions associated with these models, implying that large age gaps in these models do not necessarily correspond to extreme deviations given the observed proteomics profiles. For individuals with non-extreme standardized age gaps (i.e., normal agers), tail probabilities are generally centered near 0.5, consistent with typical regions of the conditional age distribution regardless of clock accuracy (Figure S4).

**Figure 4:**
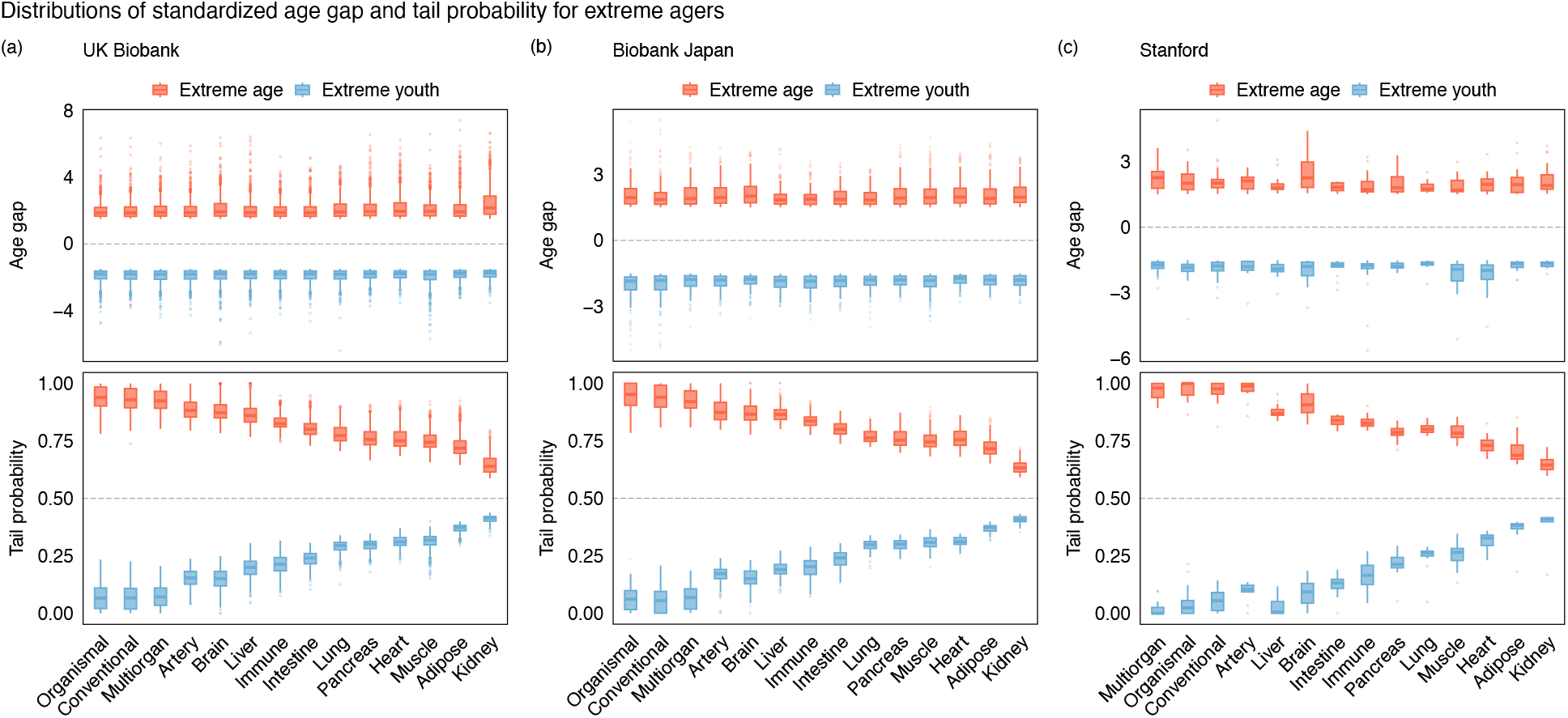
Distributions of standardized age gap and tail probability for individuals in extreme youth and extreme aging categories. Individuals are classified into extreme youth and extreme aging based on their standardized age gap (z): extreme youth (*z* < − 1.5) and extreme age (*z* > 1.5). (a) UK Biobank. (b) Biobank Japan. (c) Stanford.

These differences have important implications for the identification of extreme aging profiles at the population level. Using usual standardized age-gap thresholds, the proportion of individuals classified as exhibiting accelerated or decelerated aging was remarkably similar across clocks, despite substantial differences in predictive performance (Figure 5). In contrast, classifications based on tail probabilities (*p >* 0.9 or *p* < 0.1) varied strongly with model accuracy. Highly accurate composite clocks identified substantial numbers of individuals with strong evidence for accelerated or decelerated aging, whereas lower-accuracy organ-specific clocks identified few or virtually no individuals with comparable levels of evidence.

**Figure 5:**
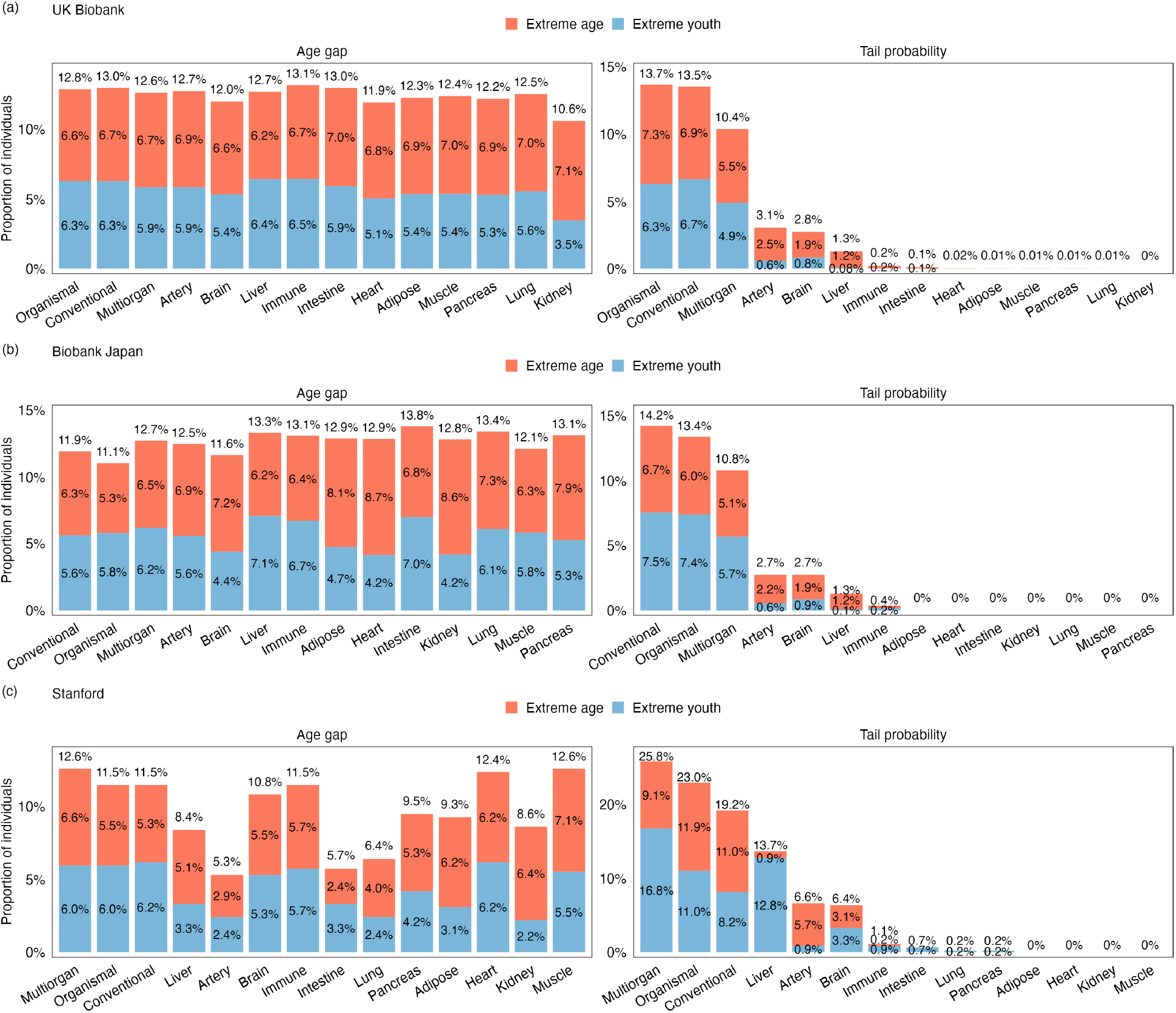
Proportion of extreme agers identified using standardized age gaps vs. tail probabilities. (a) UK Biobank. (b) Biobank Japan. (c) Stanford cohort. The left panel uses a fixed threshold (> 1.5 or < − 1.5) applied to standardized age gaps. The right panel uses a fixed threshold (*p* > 0.9 or *p* < 0.1) on the tail probability.

Overall, these findings show that reliance on standardized age-gap thresholds alone may overlook substantial model-dependent uncertainty, whereas tail probabilities provide a continuous measure of extremeness that integrates both the magnitude of the age gap and the uncertainty of the underlying prediction. As a result, individuals with similar point estimates derived from the same biological clock can exhibit differing tail probabilities if their predictive distributions vary in variance or skewness. Furthermore, individuals classified as extreme outliers based on standardized age gaps, as is common practice in the biological aging literature, may exhibit substantially different levels of predictive uncertainty, which directly affect their estimated probabilities of accelerated or decelerated aging. In particular, for the least accurate clocks, including the kidney and adipose models, prediction intervals are wide and tail probabilities remain moderate even for conventionally identified outliers (Figure 6).

**Figure 6:**
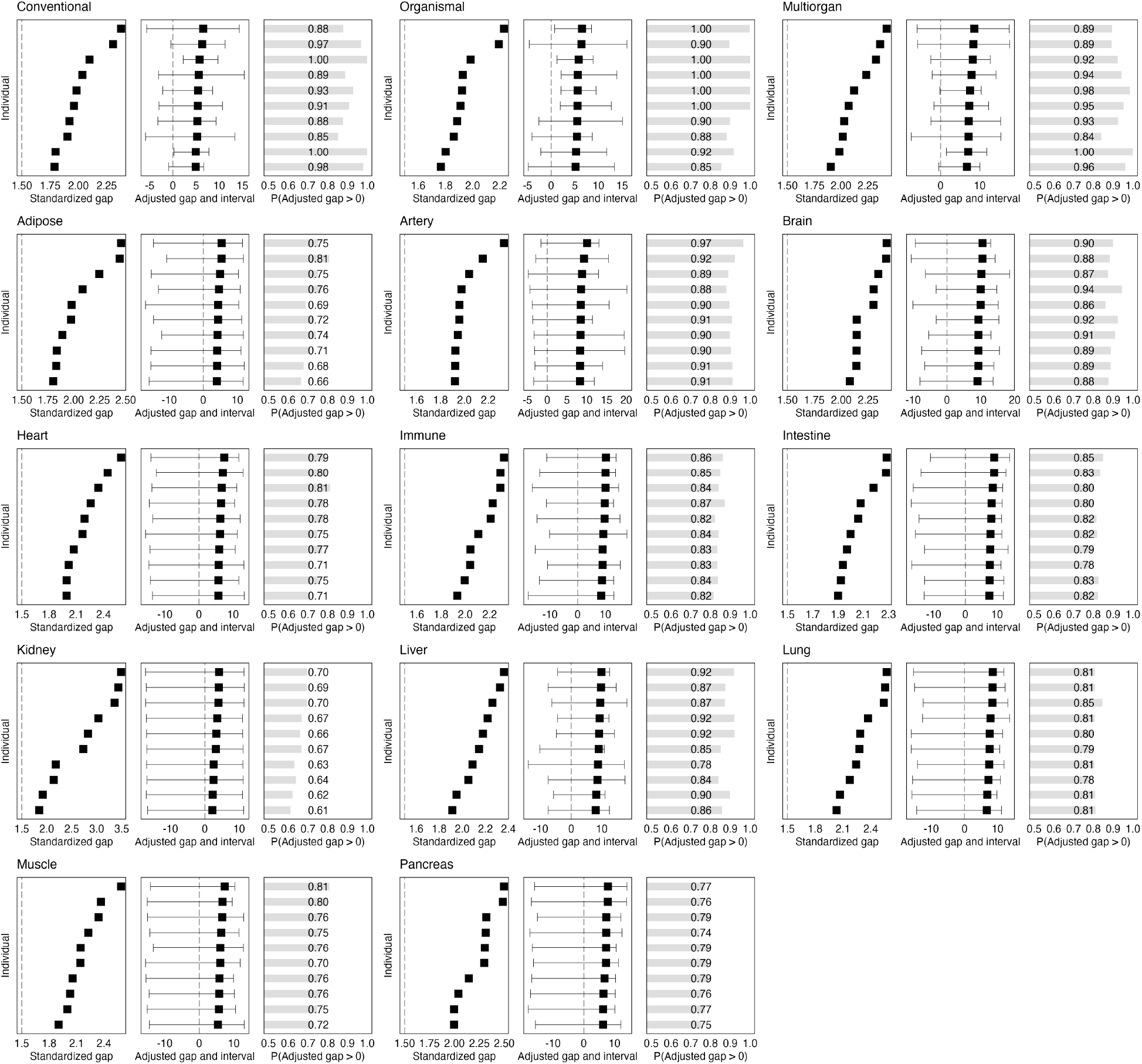
Outliers for diverse biological clocks: prediction intervals and tail probabilities in UK Biobank. For each clock, ten individuals are selected that are outliers based on the standardized age gap being greater than 1.5. For each clock, we show: Left - standardized age gaps from traditional biological clocks. Middle - Adjusted age gaps and prediction intervals from quantile regression. Right - Tail probability for adjusted age gap.

### Organ age gap, prediction interval uncertainty, mortality and disease risk

Cox proportional hazards models were employed to examine the relationship of organ-specific standardized age gaps and tail probabilities with mortality risk and risk to certain diseases (Table S2), while adjusting for chronological age and sex (Methods). To put the two predictors on the same footing, we converted standardized age gaps and tail probabilities into quintiles. Overall, both standardized age gaps and tail probability exhibited similar and significant associations with mortality and disease risk (Figure 7 for clocks trained with at least 100 proteins; Figure S5 for all clocks).

**Figure 7:**
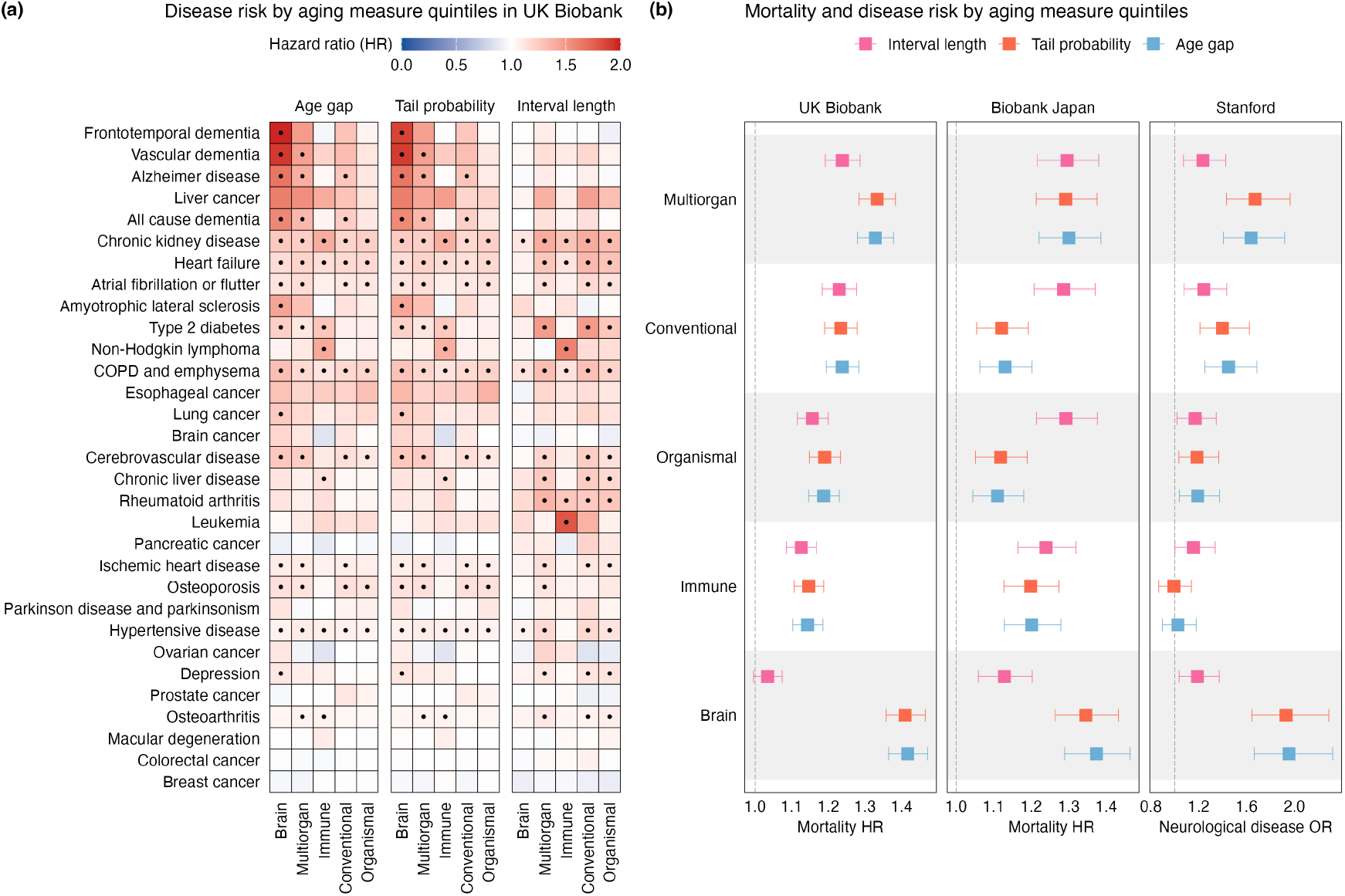
Associations of biological age measures and predictive uncertainty with mortality risk and specific diseases in UK Biobank, Biobank Japan, and Stanford cohort. (a) Disease risk associations in UK Biobank. Hazard Ratios (HR) and dots within the grid cells highlight significant associations with disease risk after Bonferroni correction. (b) Mortality (UK Biobank and Biobank Japan) and neurological disease risk (Stanford cohort) associations. Hazard Ratios (HRs) and odds ratios (ORs) along with their corresponding 95% confidence intervals are shown for various organ clocks.

In addition to biological age gaps, we examined whether predictive uncertainty itself, quantified by prediction interval length, was associated with health outcomes. We focused primarily on models trained using more than 100 proteins, as these exhibited only weak correlations between prediction interval length and age gap or tail probability (Figure S3(b–c)), suggesting that predictive uncertainty captures information largely independent of estimated biological age. Interestingly, prediction interval length was significantly and positively associated with both mortality and disease risk for these clocks (Figure 7). These findings are consistent with the hypothesis that systemic dysregulation or loss of homeostatic resilience increases the variability and unpredictability of molecular profiles, making biological age more difficult to infer accurately (Mei et al. 2023).

We also observed an opposite pattern for several organ-specific clocks, including adipose, lung, pancreas, artery, muscle, and heart, for which shorter prediction intervals were associated with greater mortality risk. These inverse associations were also observed in Biobank Japan (see replication analyses below). To better understand this behavior, we visualized the learned representations of these models using Uniform Manifold Approximation and Projection (UMAP) (McInnes et al. 2020) (Figure S6). Compared with the diffuse embeddings of the composite clocks, several organ-specific clocks exhibited pronounced one-dimensional gradients, with both tail probability and prediction interval length changing smoothly along the embedding. This observation suggests that these quantities are strongly constrained by the geometry of low-dimensional prediction models, potentially contributing to the inverse associations observed for prediction interval length.

### Replication analyses

We evaluated the reproducibility of our findings in two independent datasets: Biobank Japan and a combined clinical cohort from the Stanford Alzheimer’s Disease Research Center (ADRC) and Stanford Aging and Memory Study (SAMS). In both datasets, we applied the models trained in UK Biobank directly, without retraining, to assess their performance and generalizability under external validation (Methods).

#### Biobank Japan

We first evaluated the models in Biobank Japan, a hospital-based prospective biobank comprising approximately 267,000 participants (Nagai et al. 2017). We analyzed serum proteomic measurements from 2,962 participants generated using the Olink Explore 3072 platform, the same platform used in UK Biobank (Methods). Overall, the results were highly consistent with those observed in UK Biobank. Composite clocks showed higher *R*^2^, lower MAE, and narrower prediction intervals than organ-specific clocks (Figures S7). For composite and higher-*R*^2^ clocks, individuals with extreme standardized age scores ( |*z*| *>* 1.5) tended to have tail probabilities close to 0 or 1, indicating strong evidence for decelerated or accelerated aging, respectively. In contrast, the lower-*R*^2^ organ-specific clocks yielded tail probabilities closer to 0.5 even among individuals with extreme standardized age scores, consistent with greater predictive uncertainty (Figure 4).

Despite substantial differences in predictive performance across clocks, standardized age-gap thresholds classified similar proportions of individuals as having accelerated or decelerated aging. Tail probabilities provided a markedly different assessment: using thresholds of *p >* 0.9 or *p* < 0.1, composite clocks identified many individuals with strong probabilistic evidence of extreme aging, whereas lower-performing organ-specific clocks identified few or no individuals meeting these criteria (Figure 5).

We next examined the associations of standardized age gaps, tail probabilities, and prediction interval length with mortality using Cox proportional hazards models adjusted for chronological age and sex (Methods). The results were broadly consistent with those observed in UK Biobank. Tail probabilities and prediction interval length were positively associated with mortality for the composite clocks and for several organ-specific clocks, including the brain and immune clocks (Figure 7). We also observed inverse associations for several organ-specific clocks, similar to those observed in UK Biobank (Figure S5).

#### Stanford cohort

We next evaluated the models in a combined clinical cohort from the Stanford ADRC and Stanford SAMS. This cohort differs substantially from UK Biobank in both age distribution and ascertainment: whereas UK Biobank is a population-based cohort recruited primarily in middle age (40–69 years) with prospective follow-up, the Stanford cohort is clinically ascertained and enriched for late-life cognitive aging and dementia. To facilitate comparison with the UK Biobank training population, we therefore restricted the analysis to unique individuals aged 40-70 years with baseline proteomic measurements (*n* = 436).

The overall pattern was consistent with that observed in UK Biobank and Biobank Japan. Composite clocks achieved higher *R*^2^, lower MAE, and narrower prediction intervals than organ-specific clocks (Figure S8). Tail probabilities likewise reflected differences in predictive accuracy: for composite and higher-*R*^2^ clocks, individuals with extreme standardized age scores (|*z*|*>* 1.5) tended to have tail probabilities close to 0 or 1, whereas lower-*R*^2^ organ-specific clocks yielded probabilities closer to 0.5 even among individuals with extreme standardized age scores (Figure 4). Standardized age-gap thresholds again classified similar proportions of individuals as having accelerated or decelerated aging across clocks, despite differences in predictive performance. In contrast, tail-probability thresholds (*p >* 0.9 or *p* < 0.1) identified many individuals with strong evidence of extreme aging for the composite clocks but few or none for the lower-performing organ-specific clocks (Figure 5).

Notably, the Stanford cohort showed a higher proportion of individuals with extreme tail probabilities for the composite clocks than either UK Biobank or Biobank Japan. This pattern is consistent with the distinct clinical composition of the Stanford cohort, which includes both cognitively healthy controls and individuals with neurodegenerative disease. Relative to predictions from models trained in UK Biobank, individuals with neurodegenerative disease, particularly Alzheimer’s disease, were more likely to exhibit evidence of accelerated biological aging, whereas controls were more likely to exhibit evidence of decelerated aging, as reflected by both the distribution of tail probabilities between cases and controls and prediction interval length (Figure S9).

We further examined associations between the different biological age measures and neurodegenerative and cognitive aging disorders, including Alzheimer’s disease, mild cognitive impairment, Parkinson’s disease, and a composite neurological disease phenotype encompassing four conditions (Table S3). Dementia with Lewy bodies was excluded because of the small number of cases. Overall, associations based on tail probabilities were broadly consistent with those based on standardized age gaps, with brain, multiorgan, and other clocks showing significant associations across multiple disease outcomes (Figures 7 and S10). Despite the substantially smaller sample size, prediction-interval length was also positively associated with several neurological outcomes, including neurological disease overall, Alzheimer’s disease, and Parkinson’s disease, particularly for the composite, brain, and immune clocks. These findings provide further evidence that predictive uncertainty captures information related to disease risk beyond the magnitude of the biological age gap.

### Pipeline for uncertainty-aware biological age prediction

To enable practical implementation of the proposed framework, we developed a pipeline that takes individual-level proteomic profiles as input and automatically generates composite and organ-specific biological age estimates, together with personalized prediction intervals and tail probabilities for each of the 11 organ-specific and 3 composite clocks using models trained in the UK Biobank. The pipeline can also be used to retrain models on external datasets, enabling uncertainty-aware biological age estimation across diverse cohorts and data modalities, beyond proteomics used in this paper.

As an illustration, we present outputs for three representative individuals: one showing apparent age acceleration across multiple organs according to traditional standardized age gaps, one with age estimates consistent with normal aging, and one showing apparent age deceleration (Figure S11). These examples demonstrate how uncertainty-aware measures extend biological age assessment beyond point estimates or standardized age gaps by enabling rapid evaluation of global and organ-specific aging patterns while accounting for prediction uncertainty.

## Discussion

In conventional analyses of biological aging, outliers are typically identified using standardized age gaps, implicitly assuming that prediction uncertainty is homogeneous across individuals, such that the same age gap is considered equally informative regardless of an individual’s underlying molecular profile. It therefore does not account for the fact that some molecular profiles may yield substantially more precise age predictions than others.

Our framework instead introduces prediction uncertainty as a complementary dimension of biological aging by evaluating an individual’s age relative to the conditional distribution of age given their molecular profile. Under this perspective, the same age gap may have very different interpretations depending on the molecular features of the individual: a deviation that is highly unexpected for one molecular profile may be entirely consistent with the expected range for another. By estimating individual-specific conditional quantiles, our framework provides prediction intervals and tail probabilities that quantify the degree of discordance between chronological age and the molecular profile.

In applications to UK Biobank, we identified relatively few individuals who could be confidently classified as outliers for most organ-specific clocks. For the lower-performing clocks, individual-specific prediction intervals were often sufficiently wide that the posterior probability of accelerated or decelerated aging remained close to 0.5 for many individuals, limiting the ability to confidently identify biologically extreme individuals. A key advantage of tail probabilities over standardized age gaps is that they explicitly incorporate the full predictive distribution, thereby distinguishing between strong but uncertain deviations and weak but precise ones. In contrast, standardized age gaps provide only a point estimate and do not directly quantify this uncertainty. Consequently, standardized age gaps identify extreme individuals based solely on the magnitude of deviation relative to a global residual variance, implicitly assuming homogeneous uncertainty across individuals and not accounting for variation in predictive reliability across the proteomic feature space.

Importantly, this framework also allows prediction uncertainty itself to be studied as a measure of biological aging. Because uncertainty is estimated at the individual level, prediction-interval width captures variation in how strongly an individual’s molecular profile constrains their expected age. Thus, rather than viewing uncertainty solely as a limitation of prediction, our framework treats it as an informative property of the molecular aging landscape. Together, age gaps and their associated uncertainty provide a more nuanced characterization of biological aging by evaluating individuals relative to profile-specific uncertainty distributions.

We demonstrate that predictive uncertainty, quantified by the length of the prediction interval, is associated with mortality and disease risk beyond biological age gaps. These findings suggest that predictive uncertainty is not merely a measure of model confidence, but captures biologically meaningful variation that complements biological age gaps and provides additional information about disease risk and mortality. Longitudinal studies have shown that aging and disease are accompanied by increasing stochastic fluctuations in molecular profiles (Pyrkov et al. 2021). We hypothesize that this temporal instability leaves a cross-sectional signature, whereby individuals with greater molecular instability occupy regions of proteomic space where chronological age is inherently less predictable, resulting in wider individualized prediction intervals. Thus, prediction interval length may serve as a cross-sectional surrogate for longitudinal molecular instability and loss of physiological resilience. This interpretation is consistent with our previous finding that prediction intervals for polygenic disease risk increase with age (C. Wang, F. Wang, et al. 2025), suggesting that aging and disease may be accompanied by increasing heterogeneity in the relationships between molecular features and phenotypic outcomes.

Consistent with prior observations in the biological aging literature (Levine et al. 2018), we observed a divergence between chronological age prediction and disease risk prediction. Composite clocks trained on thousands of proteins achieved the highest accuracy for chronological age estimation, consistent with their ability to capture a broad range of age-related biological variation, while organ-specific clocks often trained on a handful of proteins achieved much lower accuracy. However, in associations with mortality and disease risk, some organ-specific clocks such as brain and kidney achieved among the strongest associations. Consequently, the strongest predictor of chronological age is not necessarily the strongest predictor of mortality. Future applications could extend this framework to models trained directly for mortality or other health outcomes, where individualized uncertainty may provide additional biological insight.

There are several important limitations of the organ-specific clocks developed in this study. First, the number of proteins available for training each organ-specific clock was relatively small, which may limit the biological resolution and predictive capacity of these models. Second, all protein measurements were obtained from plasma, and the degree to which circulating proteins capture the physiological state of individual organs is likely to vary substantially across organs. Organs that contribute more directly to the circulating proteome, such as the kidney, lung, and immune system, may be more accurately represented by plasma protein measurements than organs that are less directly reflected in circulation, such as the brain. Therefore, the observed associations between organ-specific clocks, mortality, and disease risk may partly reflect differences in the extent to which selected plasma proteins serve as proxies for the underlying biology of each organ. Future studies incorporating larger and more tissue-specific proteomic panels will be important to determine whether these differences arise from true organ-specific biology or from variation in the resolution and coverage of the available proteomic measurements.

Our replication analyses in Biobank Japan and the Stanford cohorts provide further support for our findings. Despite significant differences in the underlying cohorts, especially with the clinical Stanford data, these independent replications confirm the importance of incorporating uncertainty in biological age predictions, and suggest that tail probabilities provide a more robust characterization of individual extremity than standardized age gaps alone, particularly under conditions of cohort shift.

Collectively, this framework has important implications for personalized medicine by moving beyond point estimates of biological age deviation toward uncertainty-aware inference of biological aging. Prediction intervals and tail probabilities provide calibrated, individual-specific measures that account for heterogeneity in predictive uncertainty, enabling more reliable identification of aging profiles for which accelerated or decelerated aging is strongly supported by molecular data. By integrating both the magnitude of biological age deviation and the confidence in these estimates, this framework provides a more nuanced basis for individual-level risk stratification and precision health applications.

## Methods

### UK Biobank plasma proteomics data

The UK Biobank (UKB) is a large prospective cohort study comprising approximately 500, 000 participants recruited between 2006 and 2010 at ages 40-70 years. Extensive phenotypic and multi-omics data have been collected from the participants. The UKB Pharma Proteomics Project (UKB-PPP) consortium performed plasma proteomics profiling using the Olink Explore 3072 platform. The resulting data included measurements of 2, 923 proteins in blood plasma samples from 54, 219 UKB participants (Sun et al. 2023). Chronological age was defined as age at recruitment in the UK Biobank. Following the quality control (QC) procedures described by H. S.-H. Oh et al. 2025, samples with more than 1,000 missing protein measurements and proteins with a missingness rate exceeding 10% were excluded. After QC, the resulting dataset consisted of 44, 498 individuals and 2, 916 proteins. Missing protein abundance values were imputed using *k*-nearest neighbors (KNN) imputation in the previous study (H. S.-H. Oh et al. 2025). The samples were partitioned into training (50%), validation (10%), and test (40%) sets. Specifically, individuals from 11 randomly selected assessment centers were selected for model training (*n* = 23, 140). The remaining participants were randomly divided into a validation set to optimize the model parameters (*n* = 3, 559) and a test set to evaluate the performance of age prediction (*n* = 17, 799). To ensure consistent scaling across datasets, protein abundance values were standardized using the mean (*µ*^UKB^) and standard deviation (*σ*^UKB^) estimated from the training set, with the same transformation subsequently applied to the validation and test sets.

### Replication using Biobank Japan plasma proteomics data

We first performed replication analyses in Biobank Japan. BioBank Japan (BBJ) is a hospital-based prospective biobank comprising approximately 267,000 participants Nagai et al. 2017. All participants were diagnosed with at least one of the target diseases at participating hospitals and provided written informed consent. The study was approved by the ethics committees of the Institute of Medical Science, The University of Tokyo, and the RIKEN Center for Integrative Medical Sciences.

We analyzed serum proteomic data from 2,962 participants measured using the Olink Explore 3072 platform, same as in UK Biobank. Protein abundances were bridge-normalized across measurement batches and expressed as normalized protein expression (NPX) values using Olink NPX Explore v1.7.1. These participants were recruited between 2003 and 2008 and had been diagnosed with at least one of the following diseases: gastric cancer, colorectal cancer, breast cancer, prostate cancer, myocardial infarction, or drug eruption (https://humandbs.dbcls.jp/en/dataset/JGAD000927). Participants aged < 18 or *>* 85 years were excluded. For each protein, measurements lying more than three times the interquartile range below the lower quartile or above the upper quartile, or more than three standard deviations from the mean, were excluded. After quality control, data from 2,886 participants were included.

### Replication using Stanford plasma proteomics data

Additional independent replication analyses were performed using a pooled Stanford cohort comprising 1,636 plasma samples from the Stanford Alzheimer’s Disease Research Center (ADRC) and the Stanford Aging and Memory Study (SAMS). The ADRC and SAMS are longitudinal studies of dementia and aging that primarily enrolled older adults than those included in the UKB. Plasma proteomics profiling was performed using the Olink Explore 3072 platform that was consistent with the UKB discovery analyses, maximizing the overlap of measured proteins between the two cohorts (H. S.-H. Oh et al. 2025). To match the characteristics of the UKB discovery cohort, only baseline plasma samples from participants aged 40-70 years at sample collection were included in the replication analyses (*n* = 436). Organ age prediction models developed in the UKB were directly applied to the Stanford cohort without model retraining. Composite and organ-specific age gaps, tail probabilities and interval lengths were subsequently calculated from the corresponding age predictions using the same procedures as those applied in the UKB analyses. Clinical diagnoses, including Alzheimer’s disease, dementia with Lewy bodies, mild cognitive impairment, Parkinson’s disease, and other neurological diseases, were obtained from the corresponding Stanford cohort records. These were used to evaluate the disease associations with organ aging measures (Table S3).

The Olink proteomics data from the UKB and Stanford cohort were measured independently in different laboratories and underwent cohort-specific processing and normalization procedures. Consequently, systematic cohort-specific offsets in protein abundance were observed. Compared with the UKB, the Stanford cohort showed lower average protein abundance and greater within-protein variability. This could introduce systematic bias when directly applying the UKB-trained proteomic age models.

To reduce cohort-specific expression offsets while accounting for differences in age distributions, protein-level age-adjusted intercept calibration was performed before age prediction. For each protein, protein abundance (*x*) was regressed on chronological age (*y*) using a linear regression model, *x* = *α*_0_ + *α*_1_*y* + *ε*, separately within the UKB training set (*n* = 23, 140) and healthy controls from the Stanford cohort (*n* = 250). The fitted intercepts 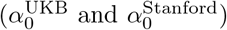 represented age-adjusted cohort-specific expression offsets.

Protein abundance for the *i*-th individual in the Stanford cohort was calibrated as 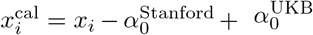. The calibrated protein abundances were subsequently standardized using 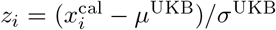, where *µ*^UKB^ and *σ*^UKB^ denote the protein-specific mean and standard deviation estimated from the UKB training set.

### Associations with mortality and disease risk

Associations between organ aging and the risk of future disease incidence or all-cause mortality were evaluated using Cox proportional hazards regression models adjusted for chronological age and sex. Disease phenotypes were defined using the International Classification of Diseases (ICD-10 and ICD-9) diagnosis codes as Argentieri et al. 2024 and H. S.-H. Oh et al. 2025, together with the corresponding diagnosis dates from the UKB. All-cause mortality with the recorded date of death were obtained from death registry records linked to the UKB. For each disease, incident cases were defined as individuals receiving a first diagnosis after baseline during follow-up. Participants with diagnoses before the baseline visit were considered prevalent cases and were excluded from the disease risk analyses. Participants who remained disease free were censored at the last available follow-up date. Followup time ranged from 0 to 18 years, with a median follow-up of approximately 10 years. For mortality risk, deaths occurring after the baseline visit were considered incident events. Participants alive at the end of follow-up were censored at the last available follow-up date (Table S2).

Separate Cox proportional hazards models were fitted for each organ age gap, tail probability, and prediction interval length adjusting for chronological age and sex. We additionally categorized each organ aging measure into quintiles within each organ model. This rank-based analysis enabled comparison of disease or mortality risk across distributional strata of aging measures and improved interpretability for effect sizes. Hazard ratio (HR), 95% confidence interval (CI), and *p* values were computed in order to quantify the association between organ aging measures (age gaps, tail probability, interval length) and the risk of incident disease or all-cause mortality. Statistical significance was determined using a Bonferroni correction across all disease outcomes, organ models, and aging measures to account for multiple testing.

In analyses of BBJ data, survival data were available for 2,878 participants Hirata et al. 2017. At baseline, the median age was 54.4 years (interquartile range [IQR], 46.0–62.1), and 2,147 (74.6%) participants were male. During a median follow-up of 11.0 years (IQR, 9.6–12.2), 535 participants died. Associations between protein abundance and overall survival were evaluated using Cox proportional hazards regression models adjusted for age and sex.

In replication analyses using the Stanford cohort, associations between organ aging measures and prevalent neurological disease status were evaluated using logistic regression, adjusting for chronological age and sex. Separate models were fitted for Alzheimer’s disease, mild cognitive impairment, and Parkinson’s disease. In addition, a composite neurological disease phenotype was defined by combining participants diagnosed with any of these four conditions and used for association tests with the same logistic regression framework. Similar to the UKB analyses, analyses based on quintiles of each aging measure were performed to facilitate fair comparison of effect sizes across the different aging measures.

### Biological age prediction and uncertainty quantification via prediction intervals and tail prob-abilities

To obtain median predictions and construct prediction intervals, we employ FS-QRPPA (Wu et al. 2026), an efficient penalized quantile regression algorithm for high-dimensional settings. Given observations (***x***_1_, *y*_1_), …, (***x***_*n*_, *y*_*n*_), with ***x***_*i*_ ∈ ℝ^*p*^ denoting the proteomic features, *y*_*i*_ ∈ ℝdenoting chronological age, and *τ* denoting the quantile level, FS-QRPPA enables parallel computation for solving the following minimization problem:

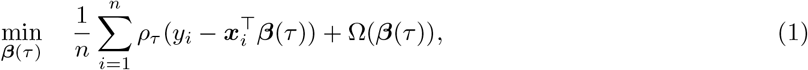

where *ρ*_*τ*_ (*z*) = (*τ* − **1** *{z* < 0*}*)*z* is the standard pinball loss for quantile regression, and Ω(·) is a piecewise linear-quadratic penalty function (Rockafellar et al. 1998). In this work, we take Ω(·) to be the elastic-net penalty.

Denote the solution of (1) by 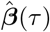. We thus obtain estimates

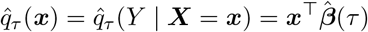

over a grid of quantile levels {*τ*_1_, *τ*_2_, …, *τ*_*M*_}, where 0 < *τ*_1_ < *τ*_2_ < … < *τ*_*M*_ < 1.

To ensure monotonicity of the predicted quantiles, we apply a common post-hoc rearrangement step by sorting the estimated conditional age quantiles (Chernozhukov et al. 2009; Chernozhukov et al. 2010) to obtain a monotonically non-decreasing sequence of predicted quantiles 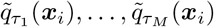. For a given 0 < *τ* < 0.5 such that *{τ*, 1 − *τ}* ⊂ *{τ*_1_, *τ*_2_, …, *τ*_*M*_ *}*, the prediction interval for the chronological age with nominal coverage level 1 − 2*τ* is then given by

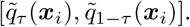

In particular, we define the biological age score for a given proteomic feature vector as the median prediction for the chronological age, 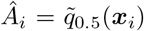. For observation *i*, let *G*_*i*_ denote the raw age gap score, which is estimated by *Ĝ*_*i*_ = *Â*_*i*_ − *y*_*i*_, where *y*_*i*_ is the observed chronological age. To reduce the systematic bias arising from the regression-to-the-mean dependence between the raw age gap and chronological age, we instead focus on the adjusted age gap 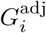, which is estimated as

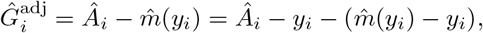

where *m*(*y*) = *E*(*Â* | *Y* = *y*) denotes the expected predicted age among individuals with chronological age *y*, and 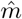 (*y*) denotes an empirical estimate of *m*(*y*). By construction,

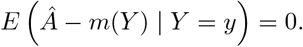

Thus, replacing *m* by 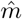 filters out the conditional mean trend of the raw age gap with chronological age, up to estimation error in 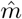.

We next construct prediction intervals with nominal coverage level 1 − 2*τ* for the adjusted predictive age gap, denoted by 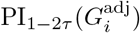, via a shift of the fitted conditional age quantile bounds by the conditional mean prediction given the chronological age *y*_*i*_:

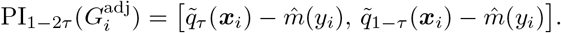

We can also report a conformalized version of 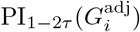 following Romano et al. (2019), which adjusts the raw quantile-regression interval to achieve marginal coverage under the exchangeability assumption.

Furthermore, we evaluate the upper-tail probability

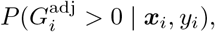

which quantifies the probability of accelerated aging relative to individuals of the same chronological age. For this purpose, we interpolate the sorted sequence of predicted quantiles 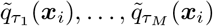 with splines that preserve monotonicity (Fritsch et al. 1980; Hyman 1983) to obtain the fitted quantile curve *Q*(*τ*; ***x***_*i*_). The predicted tail probability is given by

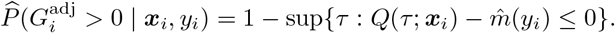

Note that 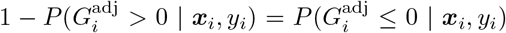 is the probability of decelerated aging relative to age-matched individuals.

## Acknowledgements

This research has been partially supported by NIH grant MH140223 and grant 2024-04735 from the Swedish Research Council (I.I.-L.). This research has been conducted using the UK Biobank Resource under Application Number 27837. Data collection was supported by the Stanford Alzheimer’s Disease Research Center, NIH/NIA grant P30 AG066515. S.N. was supported by JSPS KAKENHI (26K18279), AMED (JP223fa627001, JP24tm0424228, JP24tm0524009, JP25kk0305032, JP256f0137004), and the Japan Foundation for Applied Enzymology. Y.O. was supported by JSPS KAKENHI (25H01057), AMED (JP223fa627001, JP223fa627010, JP223fa627011, JP22zf0127008, JP23tm0524002, JP24wm0625504, JP24gm1810011, JP25kk0305022, JP26tm0135237, JP26ek0410158), JST Moonshot RD (JPMJMS2021, JPMJMS2024), the Ono Pharmaceutical Foundation for Oncology, Immunology, and Neurology, the Baelz Research Grant, and the RIKEN TRIP initiative (AGIS).

## Data Availability

The individual-level proteomics and phenotype data are available to approved researchers through the UK Biobank web portal at https://www.ukbiobank.ac.uk/. The proteome data from the Biobank Japan are available from the NBDC Human Database (https://humandbs.dbcls.jp/en/dataset/JGAD000927). The Stanford ADRC data and associated participant metadata used in this study are available upon reasonable request to the Stanford ADRC Data Release Committee (https://web.stanford.edu/group/adrc/cgi-bin/web-proj/datareq.php).

## Code Availability

Source code for the fsQRPPA package is available at https://anonymous.4open.science/r/fsQRPPA-6764/. Scripts for organ age model training, prediction, and uncertainty quantification are available at https://github.com/Iuliana-Ionita-Laza/aging-uncertainty.

## References

Argentieri, M. Austin et al. (Sept. 2024). “Proteomic aging clock predicts mortality and risk of common age-related diseases in diverse populations”. In: Nature Medicine 30.9, pp. 2450–2460. issn: 1078-8956, 1546-170X. DOI: 10.1038/s41591-024-03164-7. url: https://www.nature.com/articles/s41591-024-03164-7.

Chernozhukov, Victor, Iván Fernández-Val, and Alfred Galichon (Sept. 1, 2009). “Improving Point and Interval Estimators of Monotone Functions by Rearrangement”. In: Biometrika 96.3, pp. 559–575. issn: 0006-3444, 1464-3510. DOI: 10.1093/biomet/asp030. url: https://academic.oup.com/biomet/article-lookup/doi/10.1093/biomet/asp030.

Chernozhukov, Victor, Iván Fernández-Val, and Alfred Galichon (2010). “Quantile and probability curves without crossing”. In: Econometrica 78.3, pp. 1093–1125.

Cumplido-Mayoral, I., G. Sánchez-Benavides, N. Vilor-Tejedor, et al. (2025). “Neuroimaging-derived biological brain age and its associations with glial reactivity and synaptic dysfunction cerebrospinal fluid biomarkers”. In: Molecular Psychiatry 30, pp. 3718–3728. DOI: 10.1038/s41380-025-02961-x.

Fritsch, F. N. and R. E. Carlson (Apr. 1980). “Monotone Piecewise Cubic Interpolation”. In: SIAM Journal on Numerical Analysis 17.2, pp. 238–246. issn: 0036-1429, 1095-7170. DOI: 10.1137/0717021. url: http://epubs.siam.org/doi/10.1137/0717021.

Hirata, Makoto et al. (2017). “Overview of BioBank Japan follow-up data in 32 diseases”. In: Journal of Epidemiology 27, S22–S28. DOI: 10.1016/j.je.2016.12.006.

Horvath, S. (2013). “DNA methylation age of human tissues and cell types”. In: Genome Biology 14.10, R115. DOI: 10.1186/gb-2013-14-10-r115.

Horvath, S. and K. Raj (2018). “DNA methylation-based biomarkers and the epigenetic clock theory of ageing”. In: Nature Reviews Genetics 19.6, pp. 371–384. DOI: 10.1038/s41576-018-0004-3.

Hyman, James M. (1983). “Accurate Monotonicity Preserving Cubic Interpolation”. In: SIAM Journal on Scientific and Statistical Computing 4.4, pp. 645–654. doi: 10.1137/0904045. eprint: http://doi.org/10.1137/0904045. url: http://doi.org/10.1137/0904045.

Koenker, Roger and Gilbert Bassett (Jan. 1978). “Regression Quantiles”. In: Econometrica 46.1, p. 33. issn: 00129682. doi: 10.2307/1913643. url: https://www.jstor.org/stable/1913643?origin=crossref.

Lehallier, Benjamin, David Gate, Nicholas Schaum, et al. (2019). “Undulating changes in human plasma proteome profiles across the lifespan”. In: Nature Medicine 25, pp. 1843–1850. doi: 10.1038/s41591-019-0673-2.

Levine, Morgan E. et al. (2018). “An epigenetic biomarker of aging for lifespan and healthspan”. In: Aging 10.4, pp. 573–591. doi: 10.18632/aging.101414. url: http://doi.org/10.18632/aging.101414.

McInnes, Leland, John Healy, and James Melville (2020). “UMAP: Uniform Manifold Approximation and Projection for Dimension Reduction”. In: arXiv preprint arXiv:1802.03426. arXiv: 1802.03426 [stat.ML]. url: https://arxiv.org/abs/1802.03426.

Mei, X. et al. (2023). “Fail-tests of DNA methylation clocks, and development of a noise barometer for measuring epigenetic pressure of aging and disease”. In: Aging 15.17, pp. 8552–8575. doi: 10.18632/aging.205046.

Nagai, Akiko et al. (2017). “Overview of the BioBank Japan Project: Study design and profile”. In: Journal of Epidemiology 27, S2–S8. doi: 10.1016/j.je.2016.12.005.

Oh, D. H. et al. (2023). “Organ aging signatures in the human plasma proteome track health and disease”. In: Nature 624.7990, pp. 164–172. doi: 10.1038/s41586-023-06802-1.

Oh, Hamilton Se-Hwee et al. (Aug. 2025). “Plasma proteomics links brain and immune system aging with healthspan and longevity”. In: Nature Medicine 31.8, pp. 2703–2711. issn: 1078-8956, 1546-170X. doi: 10.1038/s41591-025-03798-1. url: https://www.nature.com/articles/s41591-025-03798-1.

Pyrkov, Timothy V. et al. (2021). “Longitudinal analysis of blood markers reveals progressive loss of resilience and predicts human lifespan limit”. In: Nature Communications 12, p. 2765. doi: 10.1038/s41467-021-23014-1.

Rockafellar, R. Tyrrell and Roger J. B. Wets (1998). Variational Analysis. Vol. 317. Grundlehren Der Mathematischen Wissenschaften. Berlin, Heidelberg: Springer Berlin Heidelberg. isbn: 978-3-540-62772-2 978-3-642-02431-3. doi: 10.1007/978-3-642-02431-3. (Visited on 07/30/2025).

Romano, Yaniv, Evan Patterson, and Emmanuel J. Candés (Dec. 8, 2019). “Conformalized Quantile Regression”. In: Proceedings of the 33rd International Conference on Neural Information Processing Systems. 318. Red Hook, NY, USA: Curran Associates Inc., pp. 3543–3553.

Rutledge, J., H. Oh, and T. Wyss-Coray (2022). “Measuring biological age using omics data”. In: Nature Reviews Genetics 23.12, pp. 715–727. doi: 10.1038/s41576-022-00511-7.

Sun, Benjamin B. et al. (2023). “Plasma proteomic associations with genetics and health in the UK Biobank”. In: Nature 622.7982, pp. 329–338.

Tanaka, Toshiko et al. (2018). “Plasma proteomic signature of age in healthy humans”. In: Aging Cell 17.5, e12799. doi: 10.1111/acel.12799.

Tian, Ye Emily et al. (2023). “Heterogeneous aging across multiple organ systems and prediction of chronic disease and mortality”. In: Nature Medicine 29.5, pp. 1221–1231. doi: 10.1038/s41591-023-02296-6.

Wang, Chen, Fan Wang, et al. (2025). “Adaptive prediction intervals for polygenic risk scores reveal individual variation in genetic predictability”. In: bioRxiv, p. 2025.10.13.682102. doi: 10.1101/2025.10.13.682102.

Wang, Chen, Tianying Wang, et al. (July 2024). “Genome-wide discovery for biomarkers using quantile regression at biobank scale”. In: Nature Communications 15.1, p. 6460. issn: 2041-1723. doi: 10.1038/s41467-024-50726-x. url: https://www.nature.com/articles/s41467-024-50726-x.

Wang, Fan et al. (Dec. 2025). “Computationally efficient whole-genome quantile regression at biobank scale”. In: Proceedings of the National Academy of Sciences 122.50, e2513007122. issn: 0027-8424, 1091-6490. doi: 10.1073/pnas.2513007122. url: https://pnas.org/doi/10.1073/pnas.2513007122.

Wang, Yu, Shuai Xiao, Bing Liu, et al. (2026). “Organ-specific proteomic aging clocks predict disease and longevity across diverse populations”. In: Nature Aging 6, pp. 162–180. doi: 10.1038/s43587-025-01016-8.

Wu, Hanqing, Jonas Wallin, and Iuliana Ionita-Laza (2026). “Scalable Ultra-High-Dimensional Quantile Regression with Genomic Applications”. In: arXiv preprint arXiv:2601.02826.

